# The effective founder effect in a spatially expanding population

**DOI:** 10.1101/006700

**Authors:** Benjamin Peter

## Abstract

The gradual loss of diversity associated with range expansions is a well known pattern observed in many species, and can be explained with a serial founder model. We show that under a branching process approximation, this loss in diversity is due to the difference in offspring variance between individuals at and away from the expansion front, which allows us to measure the strength of the founder effect, dependant on an effective founder size. We demonstrate that the predictions from the branching process model fit very well with Wright-Fisher forward simulations and backwards simulations under a modified Kingman coalescent, and further show that estimates of the effective founder size are robust to possibly confounding factors such as migration between subpopulations. We apply our method to a data set of *Arabidopsis thaliana*, where we find that the founder effect is about three times stronger in the Americas than in Europe, which may be attributed to the more recent, faster expansion.

## 2 Introduction

We may think of a range expansion as the spread of a species or population from a narrow, geographically restricted region to a much larger habitat. Range expansions are a common occurrence in many species and systems, and they happen on time scales that differ by orders of magnitude. Viruses and bacteria may spread across the globe in a few weeks Brockmann and Helbing (2013), invasive species are able to colonize new habitats over decades (Davis, 2009); and many species migrated into their current habitat over the last few millenia, following changing environments such as receding glaciers (Hewitt, 1999; Taberlet et al., 1998).

The population genetic theory of range expansion is based on two largely distinct models. The first model, based on the seminal papers of Fisher (1937) and Kolmogorov et al. (1937) and often called the Fisher-KPP model, is based on the diffusive spread of alleles, and has been mostly explored from a statistical mechanics viewpoint. The other model, called the serial founder model, has its roots in the empirically observed decrease in genetic diversity from an expansion origin Austerlitz et al. (1997); Hewitt (1999); Ramachandran et al. (2005).

The Fisher-KPP partial differential equation describes the deterministic change in allele frequency at a spatial location due to the individuals with a selected allele having more offspring than wild-type individuals. The model can be applied to range expansions by substituting the selected allele with presence of a species, and the wild-type allele as absence of a species(see e.g. Barton et al., 2013, for a recent review). Its solution is a travelling wave; similar to logistic growth, populations grow to some carrying capacity. This model has received more widespread attention recently due to the empirical tests by Hallatschek et al. (2007), who compared growing colonies of *E. coli* to the predictions from the Fisher-KPP equation.

However, it is also apparent that some of the predictions of the Fisher-KPP model are inconsistent with many macroscopic systems. In particular, the Fisher-KPP model predicts that local populations start with extremely small population sizes. This leads to a very high amount of genetic drift, and, for example, in the experiments of Hallatschek et al. (2007), all local genetic variability was quickly eliminated, so that no polymorphisms were shared between individuals sufficiently far from each other. This is in strong contrast to humans, for example, where an expansion out-of-Africa is well supported (e.g. Ramachandran et al., 2005), but where we find many genetic variants shared between all human populations. This was part of the motivation for the development of the serial founder model, which is typically based on a variant of a stepping stone model (Kimura, 1964) in one or two dimensions. Typically, only a small subset of populiations is colonized at the beginning of the process, but over time subsequent populations are colonized, usually by means of some founder effect. There are multiple ways a founder effect has been modelled: Austerlitz et al. (1997) and Ray et al. (2010) chose to model the founder effect by local logistic growth, where each local subpopulation grows according to the logistic equation. A simpler model, favored by DeGiorgio et al. (2011) and Slatkin and Excoffier (2012), is to model the founder effect by a temporary reduction in population size.

A complete serial founder model, including selection, founder events and migration between subpopulations, has, so far, be proven to complex to be of use for analytical research. However, a recursion approach (e.g. Austerlitz, 1997) and simulations (e.g. Klopfstein et al. 2006, Travis & Burton 2010) have been successfully applied to investigate the behavior of the model. Other alternatives are models that do not model expansions explicitely, and make additional simplifications. Perhaps the simplest model of this kind is that of a demographic expansion without any spatial component, which can be fully described by a change in the rate of coalescence Gravel et al. (2011); Li and Durbin (2011), with the assumption made that the population is panmictic throughout the expansion. This model resembles a spatially explicit expansion, when migration between demes in the latter are very large. A more sophisticated model, incorporating spatial structure, is the infinite island approximation model of Excoffier (2004). In this model, an originally small, panmictic population expands instantly into a metapopulation with a large number of subpopulations. In contrast to the demographic expansion, in this model we can compare coalescent time distribution between demes and within demes (called the mismatch distribution), which has been used for inference previously. However, this model assumes that all subpopulations are exchangable, so that there is no difference in coalescence times with individuals at a wave front, when compared to individuals in the center of a population. A further step were the models of DeGiorgio et al. (2011) and Slatkin and Excoffier (2012). DeGiorgio et al. (2011) derived coalescence times under a serial founder model, using a small bottleneck as a founder event. In the model of Slatkin and Excoffier (2012), the expansion is modelled as a spatial analog of genetic drift, where each founder event corresponds to a generation in a standard Wright-Fisher model.

So far, the theoretical models of range expansions have let to few applications that can be applied to interpret genetic data from non-model organisms. In this paper, we first develop a simple model of a range expansions based on a branching process approximation. The advantage of the simplicity of the model is that it leads to the development of an intuitive understanding of an expansion. We test the model using simulations, and discuss its limitations, and then show how it can be used for inference.

We demonstrate the utility of our approach by re-analyzing SNP data of the model plant species *Arabidopsis thaliana* (L.) Heynh. (Horton et al., 2012). *A. thaliana* is a small, annual plant, thought to be native to Europe, but introduced in North America and other locations (Jorgensen and Mauricio, 2004). The biogeograhy and population structure of *A. thaliana* has been well studied (Horton et al., 2012; Jorgensen and Mauricio, 2004; Nordborg et al., 2005) While earliest studies showed relatively little population differentiation on a global scale, genome-wide genetic data supports widespread population structure and clear genetic differentiation between populations (Horton et al., 2012; Nordborg et al., 2005). The availability of genome-wide SNP-chip data from more than a thousand individual plants from hundreds of locations make *A. thaliana* an ideal test case for the genetic signatures of range expansions. However, the status of *A. thaliana* as a human commensialist and the fact that *A. thaliana* is a selfing plant, may make the analysis more challenging.

## 3 Results

### 3.1 Overview of theoretical results

In this section, we will briefly outline our model and the main theoretical results. Details and full derivations can be found in the appendix. A schematic of the model studied is given in Figure 1. In brief, we assume a serial founder model on a one dimensional stepping stone grid, where initially only one deme is colonized. We compare the allele frequency of individuals in the same location as the origin of the population at time t, *X_t_*, with individuals at the wave front at time *t*, which we denote by 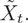 In particular, we are interested in the difference in derived allele frequency between the population at the starting position and the expansion front, which we denote as *Z_t_*. In Appendix A.1, we show that the expected difference in allele frequency is

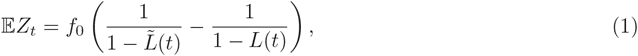

where *f*_0_ is the initial frequency of an allele, and *L*(*t*) and 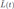 are the probabilities that an allele is lost by time *t* at the origin of the expansion and the wave front, respectively.

**Figure 1:**
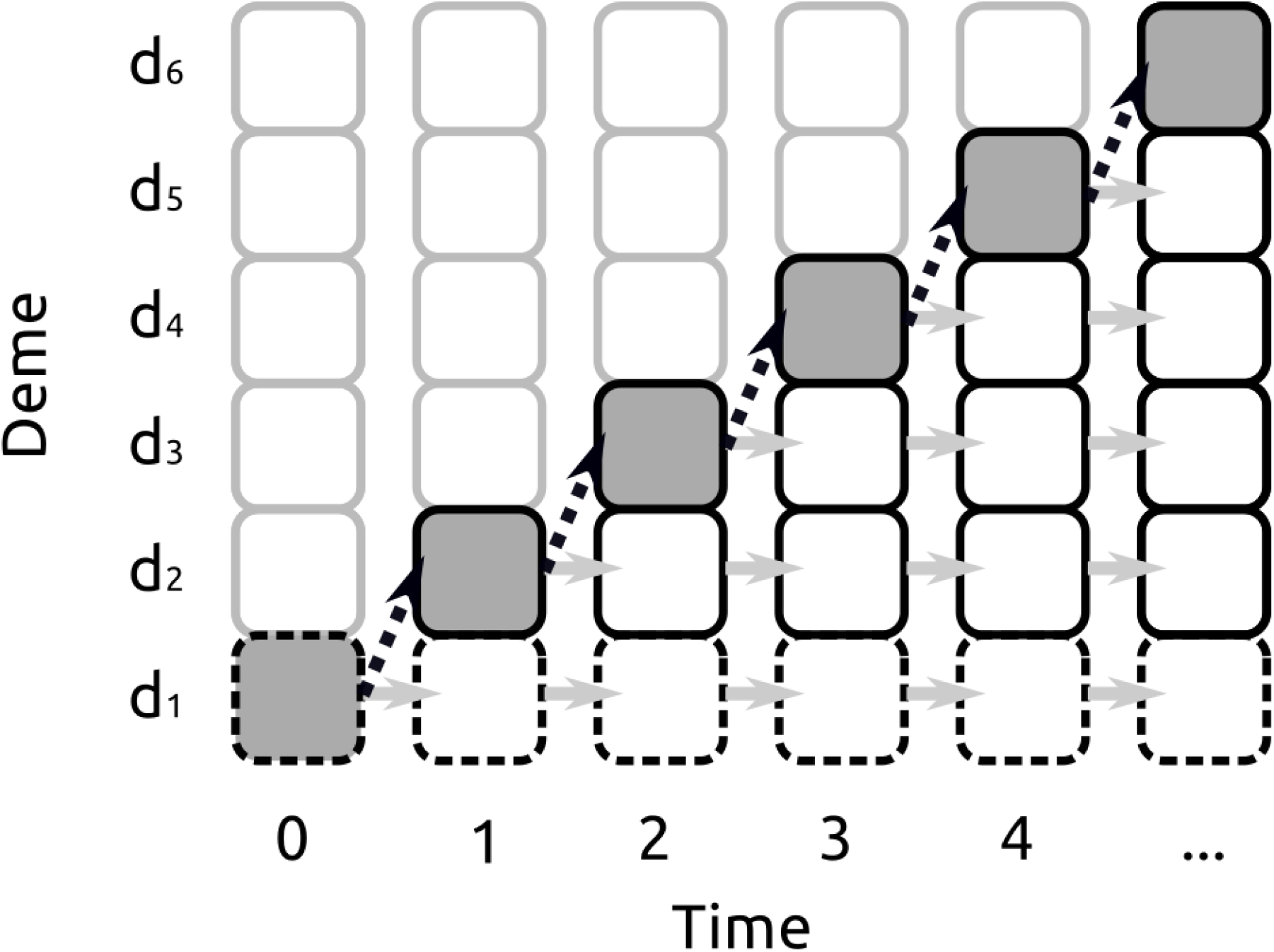
Schematic of the expansion models studied. This figure shows the basic process we study, each square corresponds to a subpopulation, with grey borders indicating subpopulations not colonized at a time step. Each time step, a new deme is colonized (black, dashed arrows), and other demes undergo neutral genetic drift (grey arrows). We compare the allele frequencies {*X_t_*} at the expansion origin *d*_1_ (dashed borderes) with the allele frequencies {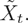} at the expansion front (dark backgrounds).

We can make this result more explicit assuming the populations evolve according to a branching process. A (Galton-Watson) branching process (Harris, 1954) models the evolution of a population by assuming that all individuals produce offspring independently from each other, with some offspring distribution *F*. In Appendix A.2 we use standard results from branching process theory to show that if each deme evolves according to a branching process, then (1) can be written as

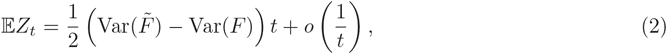

that is, the difference in allele frequency is expected to increase linearly with distance, and the slope is half the difference in the variance of offspring distribution at the expansion front 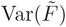 and away from the expansion front Var(*F*). Since we assume that founder effects occur at the expansion front, we expect it to have a higher offspring variance, corresponding to a lower effective population size. It is worth pointing out that the term of order *t* in 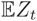 does not depend on *f*_0_, so that we expect the same slope independent of the initial allele frequency. As the higher order terms depend on *f*_0_, we will examine the accuracy of this result using simulations.

In Appendix section A.3 we then use the offspring variance from a Wright-Fisher model to define an effective founder size *k_e_*, and show tat

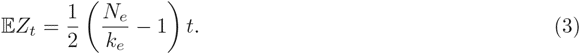

where *N_e_* is the effective population size of a deme. In some cases, it might be possible to interpret *N_e_* and *k_e_* directly. For example, if we think of a species colonizing a system of islands, *N_e_* corresponds to the carrying capacity of that species on a given island, and *k_e_* to the number of founders. In most cases, however, subpopulations are not clearly defined and the population is relatively continuously distributed. Under these circumstances, it is not clear what *N_e_* and *k_e_* represent. Therefore, we show in section A.4 that it makes more sense to think about the distance over which the ratio 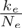 has a certain value (e.g. 0.99), that is, how far apart demes need to be so that each founding population is 1% lower than the population at equilibrium. The larger this distance, the weaker the founder effect.

Finally, in Appendix A.5 we show how we can estimate 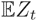 from genetic data using the *ψ* statistic defined as

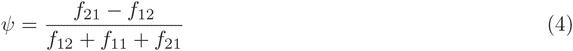

where *f_ij_* is the (*i, j*) entry in the allele frequency spectrum. *ψ* was introduced by Peter and Slatkin (2013), and we show in Appendix A.5 why *ψ* is an useful estimator of 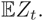 Taken together, these results suggest that we can define and estimate the effective founder effect that describes the loss of genetic diversity with distance from the expansion origin, and that we can infer the strength of the founder effect using a simple linear regression on the allele frequency of shared alleles.

### 3.2 Simulations

We validate our analytical results by performing extensive simulations under two different models. The first is the forward-in-time model stepping stone model described by Slatkin and Excoffier (2012). The second model is a backward-in-time stepping stone model, based on the Kingman coalescent (Wakeley, 2009).

#### 3.2.1 Forward simulations

We first validate our results using a discrete-time, forward-in-time Wright-Fisher model. In Figure 2, we give results for various initial allele frequencies *f*_0_, setting *k_e_* = 0.1*N* (first row), *k_e_* = 0.5*N* (middle row) and *k_e_* = 0.9*N* (bottom row). Using equation 3, we would predict *Z_t_* to be 4.5*t*, 0.5*t* and *t/*18, respectively. Those predictions are given by the red lines; the points represent data observed in simulations. We find that we get better estimates when i) the effective founder size is low, ii) the time after the expansion is low and iii) the effective population size is high. In particular, we find that we get a systematic bias when we have a very strong founder effect, and thus allele frequency differences are expected to be very large. In that case, many alleles will become fixed in the population, and the predictions between the Wright-Fisher and the branching process models are quite different.

**Figure 2:**
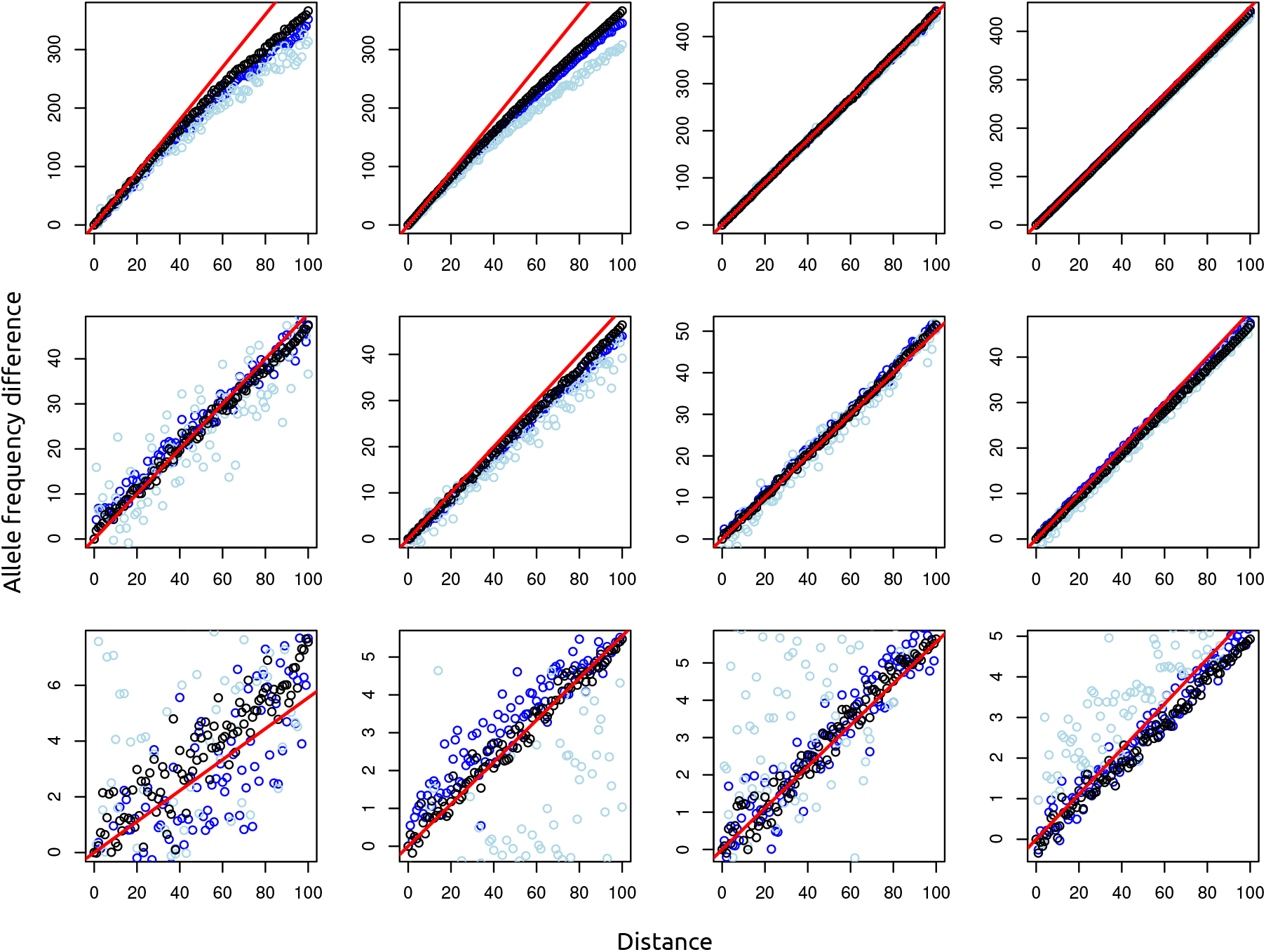
This figure shows the expected allele frequency difference between demes compared with simulations. Note: Axes labels aren’t done yet. X-axis: Deme; Y-axis: Allele count difference to deme 1. Top row: *k* = 0.1*N*, middle row: *k* = 0.5*N*, bottom row: *k* = 0.9*N*. first column: *f*_0_ = 1*, N* = 1000, second column: *f*_0_ = 10*, N* = 1000, third column: *f*_0_ = 10*, N* = 10000, Fourth column: *f*_0_ = 100*, N* = 10000, red line: prediction using branching process model. black, blue and lightblue dots correspond to samples right after expansion reached deme 100, 100 generations later and 500 generations later, respectively. Other parameters are *t* = 2, *m* = 0 and 10^6^ alleles were generated

In Figure 3, we investigate the effect of demes growing to their carrying capacity via logistic growth, as opposed to instantaneous growth which we assume in most other cases. Here we can apply the result that under non-constant founder population sizes, the effective founder size is simply the harmonic mean of all founder sizes, divided by the number of generations. In Figure 3, simulations were performed with a carrying capacity of 10,000, and growth starting

**Figure 3:**
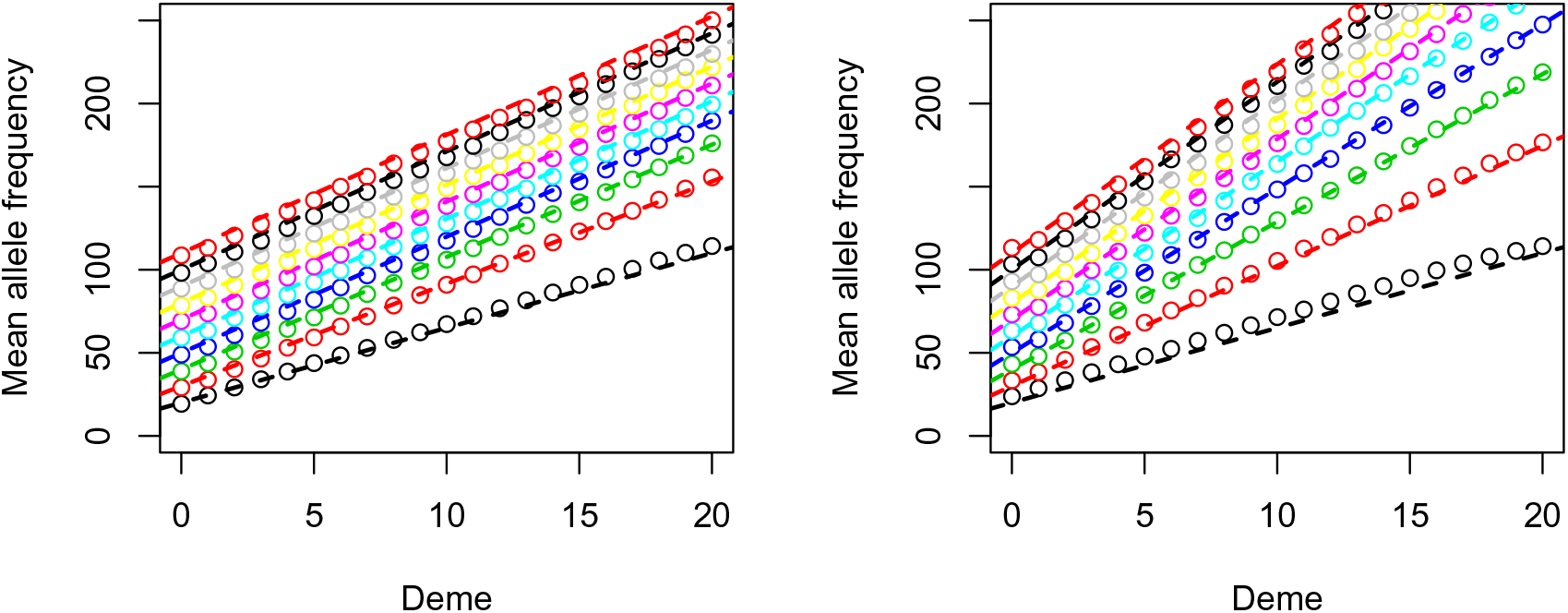
Comparison between WF-simulations and predictions from the branching process model under a logistic growth model. Growth rates were set to 1 (Panel a) and 0.5 (Panel b), respectively the lines correspond to 1-10 generations of logistic growth per expansion step (from bottom to top). Dots correspond to the simulated data, and the dashed lines are the analytical predictions using the harmonic mean of the population sizes.

#### 3.2.2 Backwards simulations

We also performed backward-in-time simulations i) to test the robustness of the branching process predictions to migration, ii) to test the effect of estimation from a subsample and iii) to to remove the initial allele frequency as an explicit parameter. Coalescent simulations were performed in a continuous-time model with discrete expansion events. In particular, most of the time lineages are allowed to merge according to the standard structured coalescent. The only exception are the expansion events, which are modelled as a single generation of Wright-Fisher mating, followed by moving all lineages in the newly colonized deme back to the founder deme. Thus, unlike the Kingman-coalescent, this model allows for multiple mergers at the wavefront. Under this model,

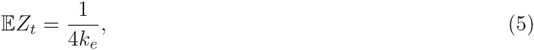

since the founder effects result in an increase of the offspring variance by a factor of (2*k_e_*)^-1^. We estimate 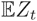 using the *ψ*-statistic defined in (Peter and Slatkin, 2013), justification of this is given in Appendix A.5. Results are displayed in Figure 4. In the top row, we show samples taken immediately after the expansion reached the boundary of the habitat, in the bottom row, we show samples that were taken a very long time (20*N* generations) after the expansion finished. We find that recent expansions are detected rather easily, almost independent of the migration rate, and the effective founder size is estimated with high accuracy. In the bottom row, we observe that for intermediate migration rates (*M* = 0.1 and *M* = 1), we still get a relatively accurate result, however, we have more noise, indicating that much larger samples would be required to obtain confident estimates, since most SNP will be either fixed or lost after this time. For a low migration rate, we see that we do not have any power for inference, since individuals all coalesce within their demes before they have the opportunity to coalesce with lineages from other locations. For high migration rates (*M* = 100), we find that the signal of the range expansion has almost vanished. Under these conditions, migration is so strong that the population essentially resembles an equilibrium isolation-by-distance population, and the signal of the range expansion has been lost. For *M* = 10, we see an intermediate behaviour, close to the origin we are at equilibrium, but far away the slope of the curve is still the same as we would expect under an expansion.

**Figure 4:**
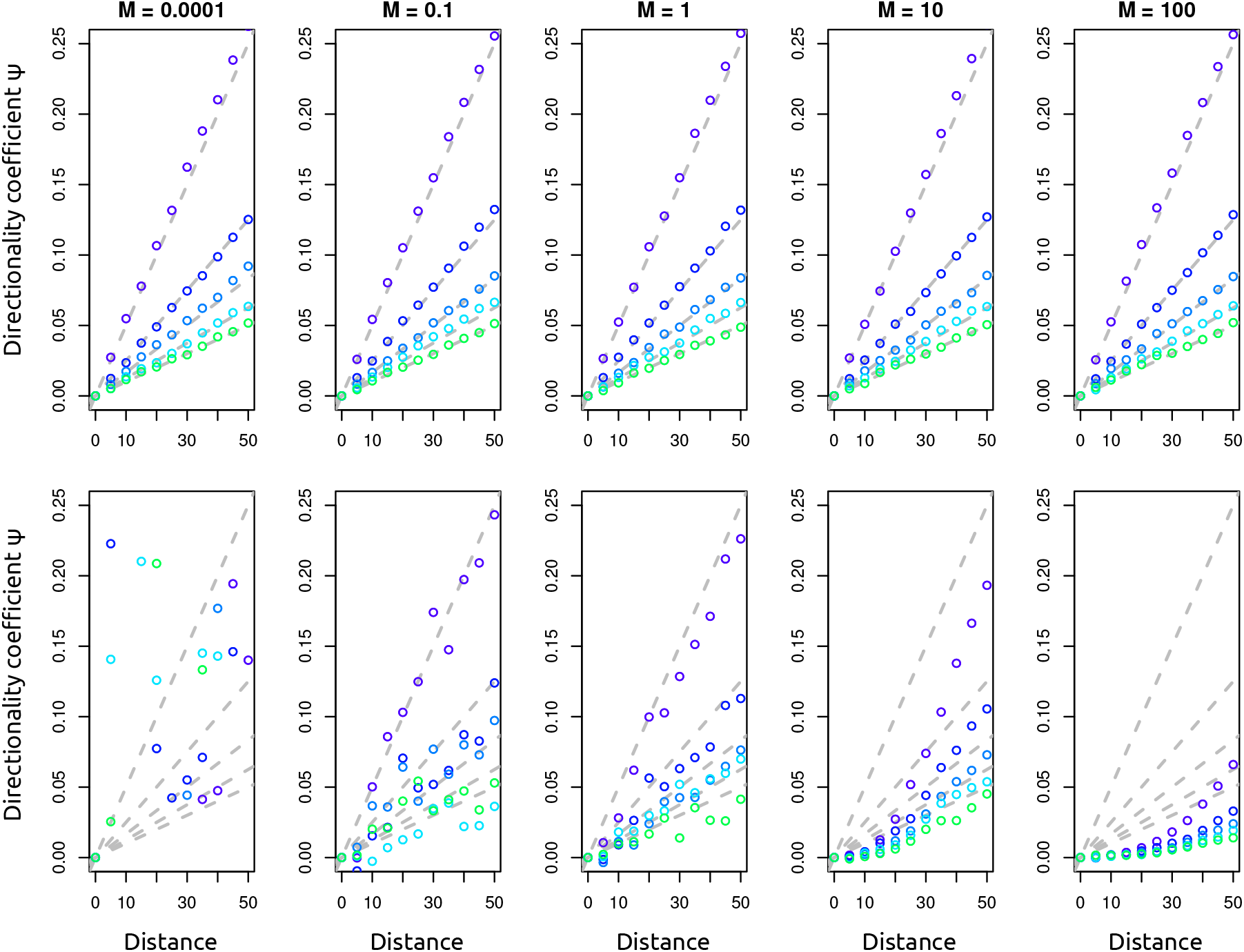
Effect of migration rate and subsampling. Each set of points corresponds to *ψ* estimated from simulations under a specific *k_e_* value, *k_e_* varies from 100 to 500 in increments of 100 (top to bottom/ blue to green). Grey dashed lines give the expectation from the branching process model. Top row: data sampled immediately after the expansion finished. Bottom row: data sampled a very long time (100 coalescence unitalics) after the expansion finished. Other parameters are: sample size *n* = 10, time between expansion events *t_e_* = 0.0001 (in coalescence unitalics).

#### 3.2.3 2D simulations

In addition to the results presented in the previous two sections, where we performed simulations in a one-dimensional habitat, we also performed simulations on 2D-stepping stone model to investigate the impact of multiple dimensions. We performed simulations by simulating expansions both under a migrant pool model and a propagule pool model (Slatkin and Wade, 1978). In a migrant pool model, all neighbouring populations send migrants at equal rates, whereas under the propagule pool model, one possible founder population is selected to send out a “propagule”, which colonizes the new deme. We find that if the sample axis is parallel to the orientation of the stepping-stone-grid, then the migration model does not matter, and we get the same behavior as in the 1D case. However, if we sample a diagonal we find that the results from the 1D-simulations are applicable under the propagule pool model, but not under the migrant pool model (Figure 5. The reason for that is quite simple: under the migrant pool model, there are many different paths on how a deme can be colonized, and the number of paths increases with distance from the origin, which reduces the amount of drift in a non-linear way with distance from the origin. In contrast, under the propagule pool model, always one path is chosen. We also find that under the 2D-model the signal of the expansion disappears faster. Whereas in the 1D-model at a migration rate of *M* = 1 the expansion is still detectable after 20*N* generations, we find that at the same migration rate, the population already approaches equilibrium.

**Figure 5:**
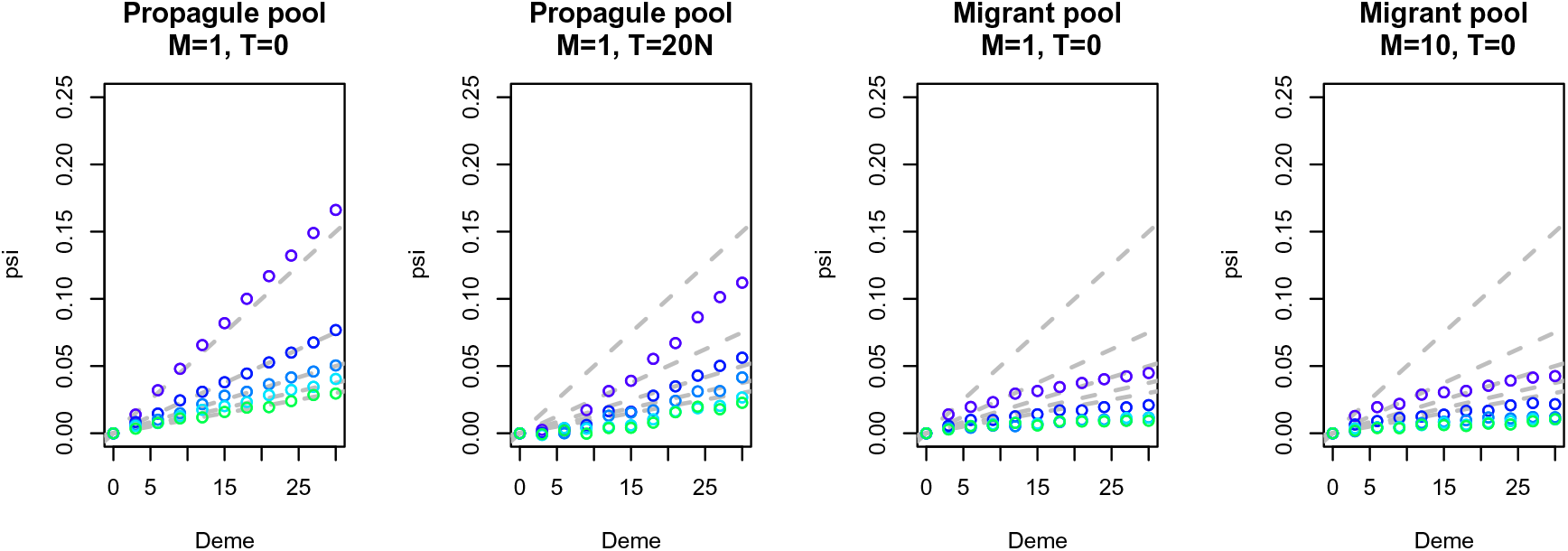
Effect of a 2D-geography. Each set of points corresponds to *ψ* estimated from simulations under a specific *k_e_* value, *k_e_* varies from 100 to 500 in increments of 100 (top to bottom/ blue to green). Grey dashed lines give the expectation from the branching process model in one dimension.

### 3.3 Application to *A. thaliana*

We applied our model to the SNP dataset of Horton et al. (2012). Based on a PCA analysis (Figure 6a) and the sample locations, we defined five regions for further analysis: Scandinavia, Americas, as well as Western, Central and Eastern Europe.

**Figure 6:**
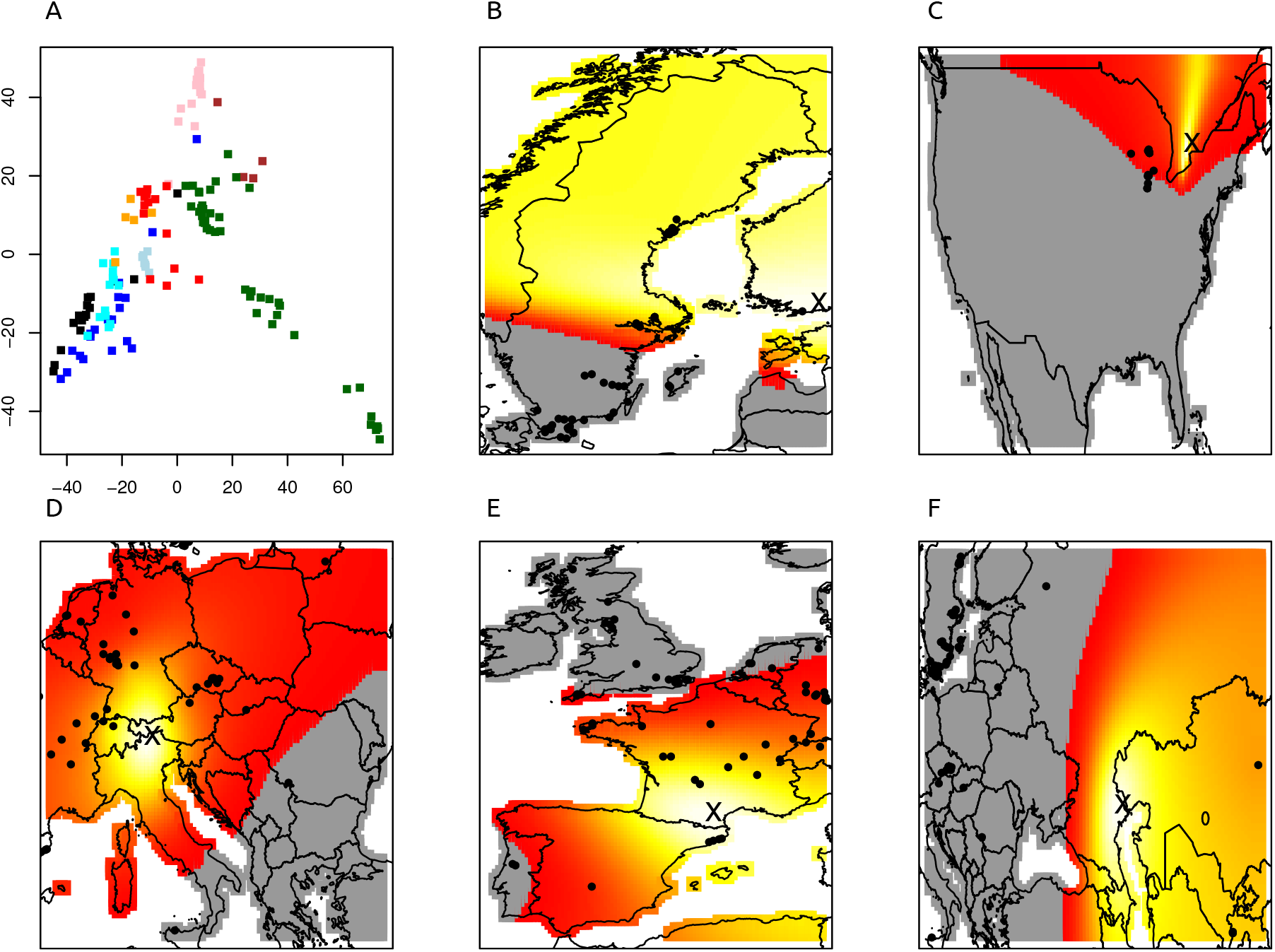
Results for *A. thaliana* data set. Panel a: PCA analysis of the 121,412 SNP. Colors: Green: Scandinavia. Black: Americas. Blue: UK. Cyan: France. Light blue: Span & Portugal. Red: North-Western Europe. Orange: Switzerland & Italy. Pink: Central Europe. Brown: Russia, Lithuania and Western Asia. Panels b-f: Expansion for Scandinavia, USA, Central Europe, Western Europe and British Isles, respectively. Brighter regions indicate more likely origin of expansion.

In the Americas, (Figure 6c) we find a most likely origin in the Great Lakes region, as opposed to the East coast. This might be somewhat surprising, but as we only have one sampling location at the East cost, power might be low. However, we can speculate that traders through the Great Lakes first introduced *A. thaliana* to the US, which may explain the signal. This may be seen as evidence for a recent introduction of *A. thaliana* into the Americas from Europe. Further support for this hypothesis comes from the fact that the American samples cluster togethr with the Western European samples in the PCA analysis, and from the fact that the North American population shows the strongest founder effect, with a 1% founder effect every 4.26 km.

Scandinavia (green dots in Figure 6a, Figure 6c), shows the most diversity within a region according to the PCA plot, and the second highest founder effect size. The most distinct samples, in the bottom left of Figure 6, are from Northern Sweden and Finland, whereas those samples that cluster with the Central and Eastern European Accessions are predominately found in Southern Sweden. We find evidence of immigration from the East, with the most likely origin of the Scandinavian accessions lying in Finland. Based on the PCA analysis, we might expect the accessions from Southern Sweden to show evidence of a range expansion from the South, and that is indeed the case when we only consider these Southern samples.

If we analyze these Southern Scandinavian samples together with the samples from Austria, Czech Republic, Russia, Lithuania and Tajikistan (Figure 6d, pink and brown dots in the PCA), we find evidence of an expansion out of eastern Asia, possibly from a refugium close to the Caspian Sea. For the Central European samples, we find an origin close to the border between Austria and Italy. This is likely a proxy for a refugium in either Southern Italy or the Balkan region, as the inferred origin was covered by an ice sheet during the last glacial maximum. Finally, for the Western European samples we find the weakest founder effect among all analyzed region, with a 1% founder effect at a scale of 38.6 km, almost an order of magnitude weaker than the strongest founder effect we observed in this set of populations, in the Americas. This is however partly due to the aggregation of the British and continental samples; if we just analyze the French, Spanish and Portugese samples (excluding the British samples), we find a founder effect of 18.7, in line with the other continental European regions. In contrast, if we analyse the British samples separately, we estimate an 1% decrease to occur over 47.8 km, and in fact we cannot exclude equilibrium isolation by distance, as, after Bonferroni correction, as*p* > 0.05.

## 4 Discussion

In this paper, we study range expansions using a serial founder model, with the main goal to develop inference procedures. We use a branching process approximation to approximate the decay of genetic diversity due to the recurring founder effects. We use this approximation to define an effective founder size, which can be estimated using standard linear regression from genetic data.

A linear or approximately linear decline of genetic diversity with distance has been observed previously in humans (DeGiorgio et al., 2009; Ramachandran et al., 2005) and in simulations (DeGiorgio et al., 2009; Peter and Slatkin, 2013). In previous work, we showed, using simulations, that the directionality index *ψ*, defined in equation (4), increases approximately linearly with distance (Peter and Slatkin, 2013). In this paper, we connect these empirical observations with a theoretical model, that explains this decay in terms of differences in offspring variance. This is justified because in populations with a higher offspring variance, genetic drift occurs faster and therefore the population’s effective size becomes smaller.

While branching processes have a long history in population genetics (Ewens, 2004), they differ from other commonly used models such as the Wright-Fisher model and the Coalescent in that the total number of individuals in the population is not constant (or following a predetermined function). Instead, the expected number of individuals is constant, leading to different dynamics. For example, a neutral branching process will eventually die out almost surely, something that cannot happen under the Wright-Fisher model. Therefore, the models presented here are only useful in parameter regions where the branching process model and other population genetic models result in similar dynamics. For example, our model breaks down if there are only few shared variants between populations. However, in this case, phylogeographic methods are arguably more appropriate than population genetic ones. Otherwise, our model appears to be useful as long as a significant fraction of variants has a most recent common ancestor during the expansion or before the expansion started. If that is not the case, as in the simulations with high *T* and high *M* in Figure 4, we find that *ψ* will be very close to zero, due to the signal of the expansion vanishing over time. The last parameter region where the model breaks down is when the mean allele frequencies become very large (Figure 2. In that case, the increase slows down due to fixations in the Wright-Fisher model, whereas it may further increase under the branching process approximation, explaining the difference.

Similar to the effective founder size defined in Slatkin and Excoffier (2012), the effective founder size *k_e_* we defined here is a variance effective size. This is different from the model proposed by DeGiorgio et al. (2009), where an explicit bottleneck was used to model a founder effect. Using an effective size is less specific than that - there are many models that will lead to the same founder size - but has the advantage that the same formalism can be applied to many different situations. We also showed various rescaling properties. Perhaps counterintuitively, 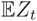 is largely independent of the expansion speed, conditional on *k_e_*. The reason for that is that, even though more segregating variants will be lost in a faster expansion, the difference between the expansion front and the rest of the population remains the same. Similarly, waiting after the expansion finished will not change 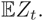

Of course, an effective size has its limitations, as in essence, it is just a measure of the speed of genetic drift. Many other models exist that may lead to similar or identical patterns of genetic diversity (DeGiorgio et al., 2009). However, in many cases it is natural to assume a range expansion occurred, often through climatic or historical evidence. In these cases, our framework may provide a starting point for a genetic analysis.

The analysis of the *A. thaliana* data shows both the usefulness and some of the limitations of our approach. We are able to identify expansion origins and infer the strength of the founder effect from genetic data. In the *A. thaliana* data set, we find that the founder effect is much stronger in the Americas than in continental Europe. This is an interesting pattern, and it would be very interesting to see if the same is true for other introduced or invasive species. In Europe, our results are consistent with previous analyses by Nordborg et al. (2005) and François et al. (2008). Nordborg et al. (2005) found that Arabidopsis likely colonized Scandinavia both from the East, through Finland, and from the South. The strong population structure is consistent with this finding a global pattern of an Eastern origin, and evidence for immigration from the South when just analyzing the Southern Swedish samples, or if we jointly analyze them with Eastern European and Asian samples. Overall, we identify a likely ice-age refugium close to the Pyrenees in Southern France or Eastern Spain, a likely refugium near the Caspian sea and a refugium in central Southern Europe, either in the Balkan or Italy, where denser sampling is required for a more accurate picture. In the Americas, we find that Arabidopsis experienced very strong founder events, and we identify a most likely point of introduction near the Great Lakes.

On the other hand, describing the founder effect as a distance over which genetic diversity decreases by a certain amount is not as satisfying as is the inference of an effective founder size, on the same scale as the effective population size. However, it is necessary because of scaling reasons; if a single population spans a larger area, then we necessarily need a strong founder effect to get the same diversity gradient than. On the other hand, if we subdivide the area of the large population into smaller populations, each of those will have its own, smaller founder effect, but the population will experience a larger number of founder events. Thus, if we know the scale of a local population, or can reasonably approxiamte it (e.g. if we know the dispersal distance of the species). We can obtain an estimate on how much lower the founder size is compared to the effective size at carrying capacity in equilibrium. On the other hand, interpreting the founder effect as a distance allows us to obtain a measure that is independent of how populations actually occupy space, which is more versatile, but somewhat harder to interpret.

## 5 Methods

### 5.1 Forward WF-simulations

Forward simulations were performed using a simple simulator implemented in R. Simulations were started with a fixed initial frequency *f*_0_, and allowed to evolve for a fixed number of generations. Every *t* - 1-th generation, the rightmost deme founded a new population, first with a single Wright-Fisher generation of size *k_e_*, which then, in the *t*-th generation, expanded to size *N*. All demes except the newly founded one underwent *t* generations of Wright-Fisher mating in the same time frame, thus after *gt* generations, *g* demes are colonized. 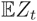 was estimated from 10^6^ replicate alleles. More complex models were implemented the same framework, i.e. we added migration between all demes at each generation and allowed the population to evolve for additional generations after the expansion finished. We also used a modification that allowed for changes in population size after each expansion event, and we used this modification to study the effect of logistic growth (see Figure 3).

### 5.2 Backwards simulations

Backward-in-time simulations were performed using the standard structured coalescent model Wakeley (2009), with a minor modification. The structured coalescent allows easy inclusion of migration events, but coalescence include migration and colonization events. The coalescent is usually studied in the continuous limit where the number of generations and population sizes are both very large. We follow this approximation with the exception of expansion events, which are modelled using a single generation of Wright-Fisher-mating. Backward in time, we stochastically merge lineages, the the backwards-transition probability for the number of lineages is (Watterson 1975, Wakeley 2008, p. 62):

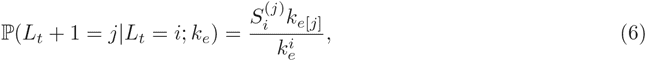

where *k_e_* is the effective founder size, *L_t_* is the number of lineages at time *t*, (time measured backwards in time in coalescence units), *S*^(*j*)^_*i*_ is the Stirling number of the second kind and *N*_[*j*]_ is the *jth* falling factorial. If the number of lineages is reduced, we merge lineages uniformly at random. All remaining lineages are then transported to a neighbouring colonized deme. To compare this model to our predictions from the branching process model, we have to consider the excess variance in offspring distribution resulting from these expansion events, which is 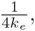 such that for this coalescent model

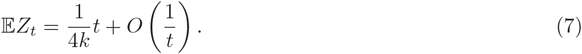

Thus, the smaller the effective founder size *k_e_*, the larger the allele frequency gradient will be. 1D- and 2D-simulations were performed using the same simulator. For 1D-simulations, we sampled eleven samples with *n* lineages every 5th deme, with 20 additional demes to avoid boundary effects. For the 2D-simulations, we sampled both a diagonal and horizontal transect. The horizontal transect, parallel to the demic structure, had length 30. The diagonal transsect, where demes were colonized every 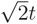 time units, had length 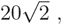 so that both transect are colonized in approximately the same time.

#### 5.2.1 Application

The data set of Horton et al. (2012), along with the coordinates for the accessions was downloaded from the project’s website at http://bergelson.uchicago.edu/regmap-data/. Genotypes of the sister species *Arabidopsis lyrata* provided by Matthew Horton were used to determine the ancestral state for each SNP. SNP where we could not determine an ancestral state unambiguously, either because no homolog *A. lyrata* allele was found, or the allele *A. lyrata* was not present in *A. thaliana*, were removed. Similarly, we removed all individuals where we did not have sampling coordinates. Since *A. thaliana* is a selfing plant and highly inbred accessions were sequenced, we only had a single haploid genotype per individual. Since our methodology requires at least two sampled haplotypes, we restricted our analysis to locations with at least two accessions sampled. To avoid bias due to very closely related accessions, we subsequently removed locations where the plants differed at less than 1.5% of sites (average heterozygosity of locations was 7.1%, with a standard deviation of 3.2%). This resulted in a total of 149 locations with at least two samples, representing 855 individuals, with 121,412 SNP genotypes remaining. As a single, uniform expansion throughout Europe seems rather unlikely, we performed a PCA analysis to find the main axes of population differentiation (Figure 6a). As the resulting pattern divided the samples broadly into four different groups, we analyzed data from these groups seperately. These groups are: Americas (black), Western Europe (blue), Central Europe (red) and Scandinavia (green). For each of these groups, we estimated the origin of the range expansion using equation 5 of Peter and Slatkin (2013). For visualization, we evaluated eq 5 of Peter and Slatkin (2013) on a grid (with locations not falling on land excluded), and estimated the best fit for the slope parameter (*v*) using linear regressions, with the location with the highest *r*^2^ corresponding to the least squared estimate of the origin of the expansion (Figure 6b-f).

The expected value of *ψ* depends on the ratio of the effective founder size *k_e_* to the effective population size *N_e_* and the number of demes that the population colonized. The number of demes is relevant, since if we subdivide the population into more demes, it will undergo more (but weaker founder effects) over the same physical distance, or conversely, if we assume that demes are large, then we have few founder events with a very strong founder effect. Using the simple model developed in this paper, we cannot distinguish these cases without additional extraneous information. For example, we may fix the size of each deme based on extraneous information. For example, if the mean dispersal distance is known for a species, we may assume that the spatial extent of each deme is approximately that dispersal distance, and we can calculate *k_e_* relative to that quantity. In this context, the ratio *k_e_/N_e_* has the interpretation as the percentwise reduction in Wright’s neighborhood size. Alternatively, if the dispersal distance is unknown, we may fix the ratio *r* = *k_e_/N_e_* to an arbitrary constant, and instead report the required distance *x_e_* over which the effective founder size is *k_e_*. This has the advantage that it provides us with a quantity that is independent of assumptions of the demic structure, and the larger *x_e_* is, the weaker is the founder effect of the population. For illustration purposes, we calculate the ratio *k_e_/N_e_* for deme sizes of 1km, 10km and 100km, as well as *x_e_* for all groups and report them in Table 1.

**Table 1.** Analysis of *A. thaliana* data.

| region | longitude | latitude | $q$ | $r_1$ | $r_{10}$ | $r_{100}$ | $x_e$ | $r^2$ | $p$ |
| --- | --- | --- | --- | --- | --- | --- | --- | --- | --- |
| Scandinavia | 24.16 | 60.32 | 0.00065 | 0.99869 | 0.9871 | 0.884 | 7.75 | 0.252 | 4.2e-55 |
| USA | -78.63 | 44.22 | 0.00118 | 0.99763 | 0.9768 | 0.808 | 4.26 | 0.242 | 6.8e-06 |
| Central Europe | 11.40 | 46.84 | 0.00035 | 0.99928 | 0.9928 | 0.932 | 14.05 | 0.171 | 8.3e-21 |
| Western Europe | 2.47 | 43.33 | 0.00013 | 0.99973 | 0.9973 | 0.974 | 38.60 | 0.264 | 4.7e-50 |
| Eastern Range | 48.29 | 46.92 | 0.00028 | 0.99943 | 0.9943 | 0.946 | 17.80 | 0.115 | 3.6e-22 |

## Acknowledgements

We would like to thank Matthew Horton for help with the data. We would also like to thank Josh Schraiber, Melinda Yang and Kelley Harris for helpful discussions. This work was supported by grant NIH-XXX to Montgomery Slatkin.

## A Derivation of main results

### A.1 Discrete time expansion model

We model a range expansion on a one-dimensional stepping stone model with potential deme positions 0, 1, 2,… labelled *d_i_, i* = 0, 1,…. All but deme *d*_0_ are not colonized at the start of the process. We denote the frequency of an allele of a biallelic marker in deme *d_i_* at time *t* as *f_i_*(*t*), and we assume that *f*_0_(0) = *f*_0_, where *f*_0_ is some constant. The population behaves as a Markov process, so that the allele frequencies at time step *t* only depend on step *t* - 1. Each time step, genetic drift will change allele frequencies according to some probability distribution. In addition, deme *d_t_* will become colonized by the offspring of individuals present at time *t*-1 in deme *d_t_*_-1_ according to some other probability distribution. For simplicity, we at first assume there is no migration between demes, and test the robustness to this assumption using simulations.

Let {*X_t_*} = {*f*_0_(0)*, f*_0_(1), … *f*_0_(*t*)} and 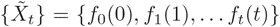 be the processes at and away from the wave front. Since we disallow migration, we can describe the history of “intermediate” demes *d_i_,* 0 < *i* < *t* by processes {*X*^(*i*)^} = {*f*_0_(0)*, f*_1_(1), … *f_i_*(*i*)*, f_i_*(*i*+1) … *f_i_*(*t*)}. In words, demes are colonized when the wave front first reaches them, and the subsequent evolution depends only on the allele frequencies at the time when they first evolved. From this construction, it follows that for *i* < *j*, {*X*^(*i*)^_*t*_} and {*X*^(*j*)^_*t*_} are conditionally independent given *f_i_*(*i*). Together with the Markov property this implies that the difference in allele frequency in two demes is a function of distance, i.e. they obey

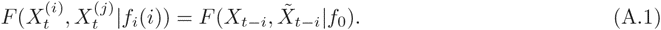

Throughout this section, we assume that 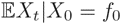 is constant, which is satisfied if there are new new mutations and no selection, and we further assume that Var(*X_t_*) < ∞. For example, for the critical branching process model we introduce in the following section, Var(*X_t_*) = *σt*, where *σ* is the offspring variance in one generation. Then the autocovariance for *s < t* is,

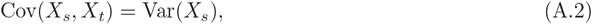

and similarly for 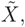 because {*X_t_*}, 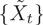 are martingales.

Next, we define the conditioned processes {*Y_t_*} = {*X_t_*|*X_t_ >* 0} and 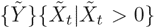 which give the allele frequency conditional on the allele not being lost.

Then, we have that

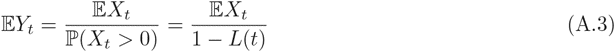

since

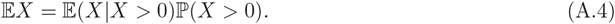

Here, 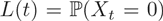 denotes the probability that an allele is at frequency zero in generation *t*, and we remove the dependency of *L*(*t*) from *f*_0_ for notational convenience.

Using the conditional variance formula, we can compute the variance and autocovariance of {*Y_t_*}:

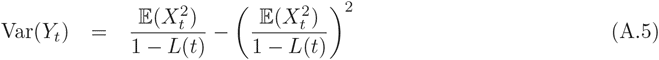

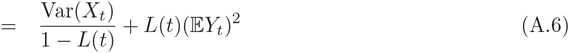

and covariance for *s* < *t*

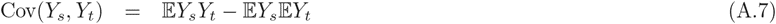

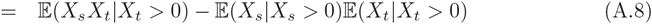

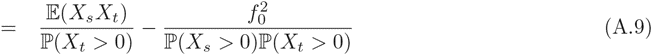

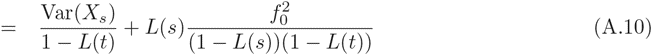

The last quantity of interest is the difference 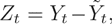 which gives the difference in allele frequency between the wavefront and the origin of the expansion, conditional on an allele surviving in both locations. We find that

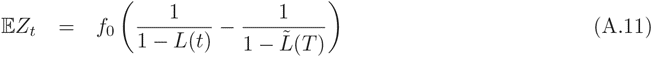

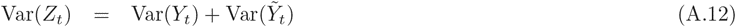

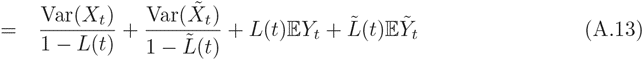

and

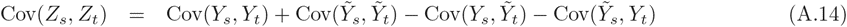

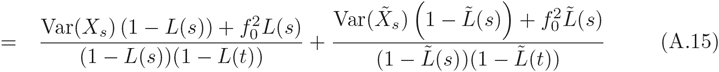

### A.2 Branching process

To further specify the moments derived in Appendix A.1, we need to define Var(*X_s_*), *L*(*s*) and *f*_0_, and the corresponding quantities at the wave front. This is particularly easy using a Galton-Watson branching process. Under this model, each generation individuals leave offspring independent from each other according to some offspring distribution *F*. Let *L_i_*(*t*) denote the probability that an allele has been lost by generation *t*, given that it started with *i* copies in generation 0. Kolmogorov (1938) showed that when *t* is large, *L*_1_ is well approximated by

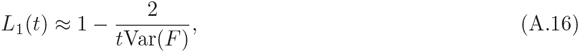

where *F* is the offspring distribution and Var*F* is assumed to be finite. We assume that a branching process with offspring distribution *F* describes neutral genetic drift at the wave front, and that the colonization of new demes occurs according to a branching process with offspring distribution 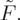

If the initial frequency *f*_0_ of the allele is greater than one, the corresponding expression becomes

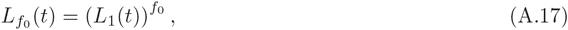

by independence of individuals. Using a Taylor expansion around *t* = ∞ yields

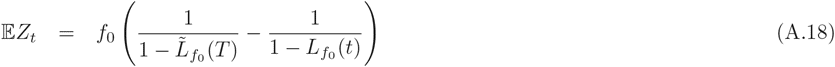

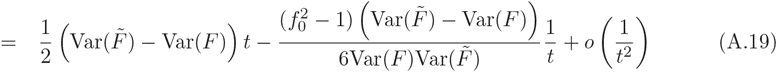

Thus, we find that the expected difference in allele frequency between the expansion origin and the front of the population increases approximately linear with distance, the slope of the curve being the difference in offspring variance of individuals at the wavefront and expansion origin. From the second term in the Taylor expansion we see that the approximation is suitable when *t > f*^2^_0_, i.e. the number of demes between the two samples is large, and the frequency of the allele at the founding location is small.

### A.3 Effective population size

The variance effective population size for a Cannings model is defined as

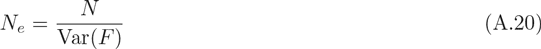

where *N* is the absolute number of individuals per population. The branching process considered above is not a Cannings model, however, the evolution of the offspring of a single individual under a Cannings population is well approximated by a branching process, as long as that offspring only makes up a small fraction of the individuals in a population. Fisher (1930) pioneered the modelling of population genetics using branching processes (Ewens, 2004, p.29). Under a Wright-Fisher model, the offspring distribution of a single individual has mean and variance very close to one. This justified Fisher to approximate the evolution of an individual under the Wright-Fisher model as evolving according to a branching process with a Poisson(1) offspring distribution, which has offspring variance 1, a model we will also use here to model genetic drift away from the wave front.

To incorporate the reduced effective size of a founder effect at the wave front, we use a modified offspring model: with probability (1-*α*), an individual at the wavefront does not produce any offspring. With probability *α*, the number of offspring is Poisson distributed with parameter 1*/α* s.t. the overall expected number of offspring is still one and the variance is 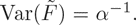 This allows us to define an effective founder size *k_e_*

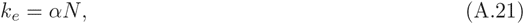

which measures the “increase” in genetic drift at the wave front.

Combining eq. A.21 and eq. A.19 yields

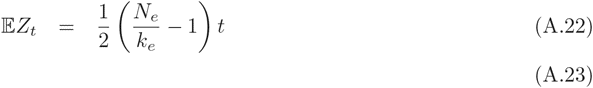

From this, we see immediately that 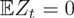 only if *N_e_* = *k_e_*, and also that the effective founder size enters the equation only in the ratio 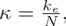 so that it makes sense to further define the relative founder size *κ*, which measures the strength of the founder effect.

### A.4 Rescaling

The branching process we used above assume that exactly one generation of genetic drift happens between each founder event. In this section, we show that the expected allele frequency difference between the expansion front and at the origin is (i) invariant of additional generations between expansion events and (ii) invariant to additional generations after the expansion finished.

Both follow from the fact that for a branching process with mean 1, the variances of subsequent generations can simply be added: Consider the generating function of a critical branching process *B* after *t* generations, denoted by *p_t_*(*s*) which has variance *p_t_*(1)^′′^. Then, after an additional generation, the generating function becomes *q*(*p_t_*(*s*)), where *q*(*s*) is the generating function of the offspring distribution of that additional generation. Then, the variance in offspring after this additional generation is *q*(*p_t_*(1))″.

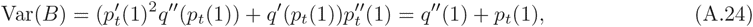

since 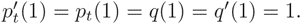

Thus, if individuals in the range expansion model have offspring variance *v* at the expansion front and variance 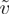 away from the front, the total variance after *t* time steps with *d* expansion events is 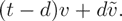

Now from eq. (A.19) we have (for *f*_0_ = 1),

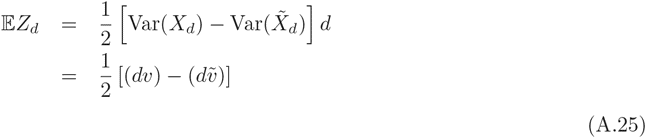

Adding *T* generations with neutral drift between each founder event and *τ* generations after the expansion stopped, changes this only to

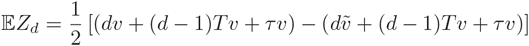

which simplifies to eq. (A.19).

We can model more complex expansion models, such as an extended bottleneck or logistic growth similarly. Again, this will result in an increase of *Var*(*X_d_*) and 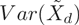 by the same amount, which cancels in the difference.

Furthermore, we can also change how we subdivide a population into demes. It is easy to see that a population with expansions at times 0, 1, 2*, …* and offspring variances Var(*F)* and 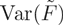 behaves similarly to a population with expansions occuring at times 0*, δt,* 2*δt, …* with offspring variances 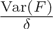 and 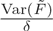 in the sense that 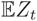 will be the same for either population. This suggests that it is not important how we subdivide space into demes, only the relative size of the founder population versus the neutral populations matters. Thus, it is most convenient to report the strength of the founder effect in units of “decrease in genetic diversity per unit of distance.

### A.5 Estimation

To estimate E*Z_t_* from genetic data, we need to take subsampling into account, i.e. we need to estimate 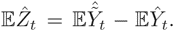 In particular, the probability that an allele got lost from a population is not the same as it being absent from a sample. To model subsampling, we assume we start with *f*_0_ copies of the derived allele and *A*_0_ copies of the ancestral allele, all evolving as a independent branching processes. The expected number of ancestral alleles will be 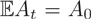 in all generations, whereas the expected number of the derived allele, conditioned on it not being lost, is 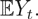 We Hence, in generation *t*, the probability of drawing *m* copies of the derived allele out of *n* samples is approximately binomially distributed with parameters *n* and 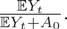 The mean of the expected allele frequency, conditional of sampling at least one derived allele is

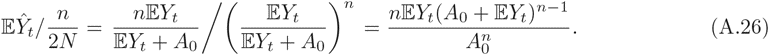

with the 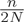 term normalizing 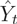 to allele counts. Setting 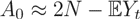 we obtain the series representation

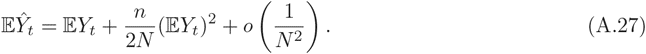

Hence,

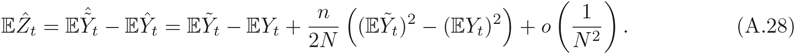

and we see that we have a bias term that increases with sample size. Hence, the easiest way to proceed is to downsample larger samples to a sample size of two, the case that is arguably most important in light of genomic data.

To compare samples of size *n*_1_ and *n*_2_, from a site frequency spectrum S = *f_ij_*, 0 ≤ *i* ≤ *n*_1_, 0 ≤ *j* ≤ *n*_2_ we can calculate a reduced site frequency spectrum matrix *S*′ from the full site frequency spectrum using

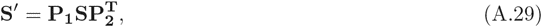

where **P**_1_ and **P**_2_ are (2 + 1) × (*n*_1_ + 1) and (2 + 1) × (*n*_2_ + 1) matrices (with indeces starting at 0), respectively, with entries

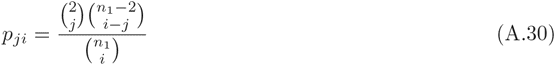

for 0 ≤ *i* ≤ *n*_1_ and 0 ≤ *j* ≤ 2 for **P**_1_. Entries in **P**_2_, are similar, except *n*_1_ is replaced by *n*_2_.

If we denote the entries of **S**′ with *s_ij_*, we can write 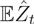 as

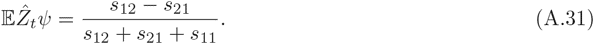

This statistic is identical to the *ψ* statistic defined in Peter and Slatkin (2013), where we did not give any theoretical justification.

The table shows the inferred latitude and longitude of the origin. *q*: regression slope in *km*^-1^, *r_i_* = *k_e_/N_e_*, for demes of size *ikm*. *d_i_*, distance (in *km*) over which 1 − *k_e_/N_e_* = 1%. *r*^2^ and *p*: adjusted coefficient of determination and Bonferroni-corrected *p*-value.

## References

Austerlitz F, Jung-Muller B, Godelle B, Gouyon PH. 1997. Evolution of coalescence times, genetic diversity and structure during colonization. Theoretical Population Biology. 51:148–164.

Barton NH, Etheridge AM, Kelleher J, Véber A. 2013. Genetic hitchhiking in spatially extended populations. Theoretical Population Biology. 87:75–89.

Brockmann D, Helbing D. 2013. The hidden geometry of complex, network-driven contagion phenomena. Science. 342:1337–1342.

Davis MA. 2009. Invasion Biology. Oxford; New York: Oxford University Press, 1 edition.

DeGiorgio M, Degnan JH, Rosenberg NA. 2011. Coalescence-Time distributions in a serial founder model of human evolutionary history. Genetics. 189:579–593. PMID: 21775469.

DeGiorgio M, Jakobsson M, Rosenberg NA. 2009. Explaining worldwide patterns of human genetic variation using a coalescent-based serial founder model of migration outward from africa. Proceedings of the National Academy of Sciences. 106:16057–16062.

Ewens WJ. 2004. Mathematical Population Genetics: Theoretical introduction. Springer.

Excoffier L. 2004. Patterns of DNA sequence diversity and genetic structure after a range expansion: lessons from the infinite-island model. Molecular Ecology. 13:853–864.

Fisher RA. 1937. The wave of advance of advantageous genes. Annals of Eugenics. 7:355–369.

François O, Blum MGB, Jakobsson M, Rosenberg NA. 2008. Demographic history of european populations of arabidopsis thaliana. PLoS Genet. 4:e1000075.

Gravel S, Henn BM, Gutenkunst RN, et al. (560 co-authors). 2011. Demographic history and rare allele sharing among human populations. Proceedings of the National Academy of Sciences. p. 201019276. PMID: 21730125.

Hallatschek O, Hersen P, Ramanathan S, Nelson DR. 2007. Genetic drift at expanding frontiers promotes gene segregation. Proceedings of the National Academy of Sciences. 104:19926–19930.

Harris TE. 1954. The theory of branching processes. Courier Dover Publications.

Hewitt GM. 1999. Post-glacial re-colonization of european biota. Biological Journal of the Linnean Society. 68:87–112.

Horton MW, Hancock AM, Huang YS, et al. (13 co-authors). 2012. Genome-wide patterns of genetic variation in worldwide arabidopsis thaliana accessions from the RegMap panel. Nature Genetics. 44:212–216.

Jorgensen S, Mauricio R. 2004. Neutral genetic variation among wild north american populations of the weedy plant arabidopsis thaliana is not geographically structured. Molecular Ecology. 13:3403–3413.

Kimura M. 1964. Diffusion models in population genetics. Journal of Applied Probability. 1:177–232.

Kolmogorov A, Petrovskii I, Piscounov N. 1937. A study of the diffusion equation with increase in the amount of substance, and its application to a biological problem. Univ., Math. Mech. 1:1–25.

Li H, Durbin R. 2011. Inference of human population history from individual whole-genome sequences. Nature.

Nordborg M, Hu TT, Ishino Y, et al. (24 co-authors). 2005. The pattern of polymorphism in arabidopsis thaliana. PLoS Biol. 3:e196.

Peter BM, Slatkin M. 2013. Detecting range expansions from genetic data. Evolution. 67:3274–3289.

Ramachandran S, Deshpande O, Roseman CC, Rosenberg NA, Feldman MW, Cavalli-Sforza LL. 2005. Support from the relationship of genetic and geographic distance in human populations for a serial founder effect originating in africa. Proceedings of the National Academy of Sciences of the United States of America. 102:15942–15947.

Ray N, Currat M, Foll M, Excoffier L. 2010. SPLATCHE2: a spatially explicit simulation framework for complex demography, genetic admixture and recombination. Bioinformatics. 26:2993–2994.

Slatkin M, Excoffier L. 2012. Serial founder effects during range expansion: a spatial analog of genetic drift. Genetics. 191:171–181. PMID: 22367031.

Slatkin M, Wade MJ. 1978. Group selection on a quantitative character. Proceedings of the National Academy of Sciences of the United States of America. 75:3531–3534. PMID: 16592546 PMCID: PMC392812.

Taberlet P, Fumagalli L, Wust-Saucy A, Cosson J. 1998. Comparative phylogeography and postglacial colonization routes in europe. Molecular Ecology. 7:453–464.

Wakeley J. 2009. Coalescent theory: an introduction. Roberts & Co. Publishers.

